# The identity and distribution of *Bhavania annandalei* Hora 1920 (Cypriniformes: Balitoridae), a hillstream loach endemic to the Western Ghats of India

**DOI:** 10.1101/2020.04.24.060103

**Authors:** Remya L. Sundar, V.K. Anoop, Arya Sidharthan, Neelesh Dahanukar, Rajeev Raghavan

## Abstract

*Bhavania annandalei* Hora 1920, is resurrected from the synonymy of *B. australis* (Jerdon 1849) based on examination of freshly collected topotypic specimens. The two species can be distinguished by a combination of morphological, morphometric and meristic characters, and by genetic distance using mitochondrial *cox1* gene. The distribution of *B. annandalei* is restricted to the river systems draining the Agasthyamalai hills, below the Shencottah Gap in southern Western Ghats.

---

The Hillstream loach *Bhavania annandalei*, was described by Hora (1920; p203) from Tenmalai, erstwhile Travancore State (= current day Southern Kerala), and suggested that the species occurs throughout the southern Western Ghats in the Nilgiris, Malabar and Travancore. Hora (1920) diagnosed *B. annandalei* from its only known congener, *B. australis* (Jerdon 1849) (type locality: Walliar Jungle = Walayar), by a combination of characters the most prominent of which included a broad snout (vs. pointed), interrupted lower lip (vs. continuous), caudal-lobes equal (vs. lower lobe longer), and presence of a pair of papillae on the lower lip (vs. absence).

Hora’s (1920) description of *B. annandalei* was however, based on a single adult female specimen collected by Dr. Annandale from Travancore. Though Hora (1920) seemed to have access to additional juvenile specimens collected by Captain Sewell from the Nilgiris (Cherambadi) and Wayanad (Nellimunda, Mananthavady and near Vythiri), he did not examine them or provide other details. Subsequently, Hora (1937; p8) extended the distribution of the species to Mysore, based on four specimens collected by M.S Bhimachar from a stream between Kottigehar and Balehonnur (erstwhile Mysore State = current day Tunga River System in Karnataka). No details of the specimens were provided.

In his review on ‘Homalopterid fishes from Peninsular India’, Hora (1941) synonymized *B. annandalei* with *B. australis*, after examining specimens from throughout its distribution range including Kallar/South Travancore (current day Vamanapuram River, Kerala); Pampadumpara/North Travancore (current day Periyar River, Kerala); Sethumadai Hills/ Mysore (current day Anamalai hills near Pollachi, Tamil Nadu); and Kottigehar/Mysore (current day Tunga River, Karnataka), and realizing that his description of *B. annandalei* was based mainly on immature specimens. This synonymy was subsequently adopted by Menon (1987) in his review of the homalopterid loaches of India, but without examining the type (or fresh topotypes) of *B. annandalei*, or the topotypes of *B. australis*. Later workers followed this synonymy and considered *Bhavania* to be monotypic (Talwar & Jhingran 1991; Menon 1999, Kottelat 2012).

Given their hill-stream adaptations (Hora 1920, 1937, 1941), and the fact that the type locality of *B. annandalei* (Tenmalai) and *B. australis* (Walayar) are at least 300km apart and separated by two significant biogeographic barriers - the Palghat Gap and the Shencottah Gap (see Anoop et al. 2018), it is highly unlikely that the two are conspecific. Collection of fresh topotypic specimens of both *B. australis* and *B. annandalei* and detailed examination and comparison of their biometrics, and genetic distance analysis based on the mitochondrial *cox1* gene, revealed that the two species are clearly distinct. We therefore resurrect *Bhavania annandalei* Hora 1920, from the synonymy of *B. australis* (Jerdon 1849) and provide notes on the distribution range of this species.

Six specimens of putative topotypic *Bhavania annandalei* were collected from Palaruvi falls at Tenmala (Kallada River), Kerala, and six specimens of putative topotypic *B. australis* were collected from near the Kavarakund falls, upstream of Malampuzha Reservoir, Kerala, India (Fig. 1). Specimens collected in the current study are in the museum collection of the Kerala University of Fisheries and Ocean Studies (KUFOS), Kochi, India. Morphometric measurements were taken for 37 characters (measured to the nearest 0.1mm using digital callipers) and meristic values were determined for 10 characters using a stereo-zoom microscope (Table 1 and 2). For statistical analysis of morphometric data, subunits of body were taken as percentage of standard length and subunits of head were taken as percentage of head length. Principal component analysis (PCA) was performed to check whether the two species formed distinct clusters in multivariate space using correlation matrix. Null hypothesis that the clusters are not significantly different from each other was tested using Analysis of similarities (ANOSIM) employing Euclidian distances and 9999 permutations. Statistical analysis was performed in PAST 4.02 (Hammer et al. 2001). Genetic sequences of mitochondrial partial cytochrome oxidase subunit 1 (*cox1*) were obtained from our related study (Sidharthan et al., In Review). Gene sequences were aligned using MUSCLE (Edgar 2004) and raw genetic distance was estimated using MEGA 7 (Kumar et al. 2016).

**Figure 1.**
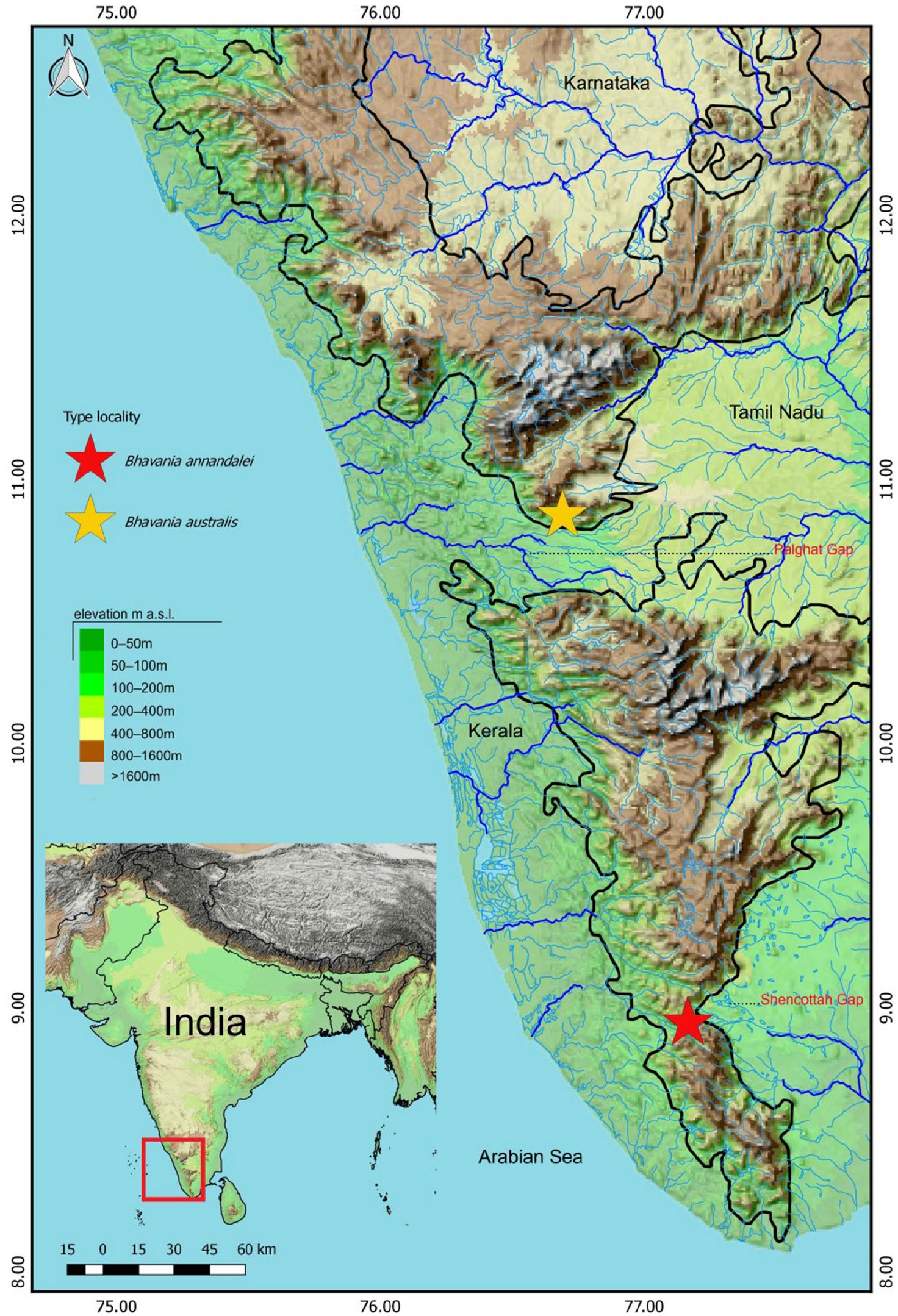
Collection localities of putative topotyes of *Bhavania annandalei* and *B. australis*.

**TABLE 1.**
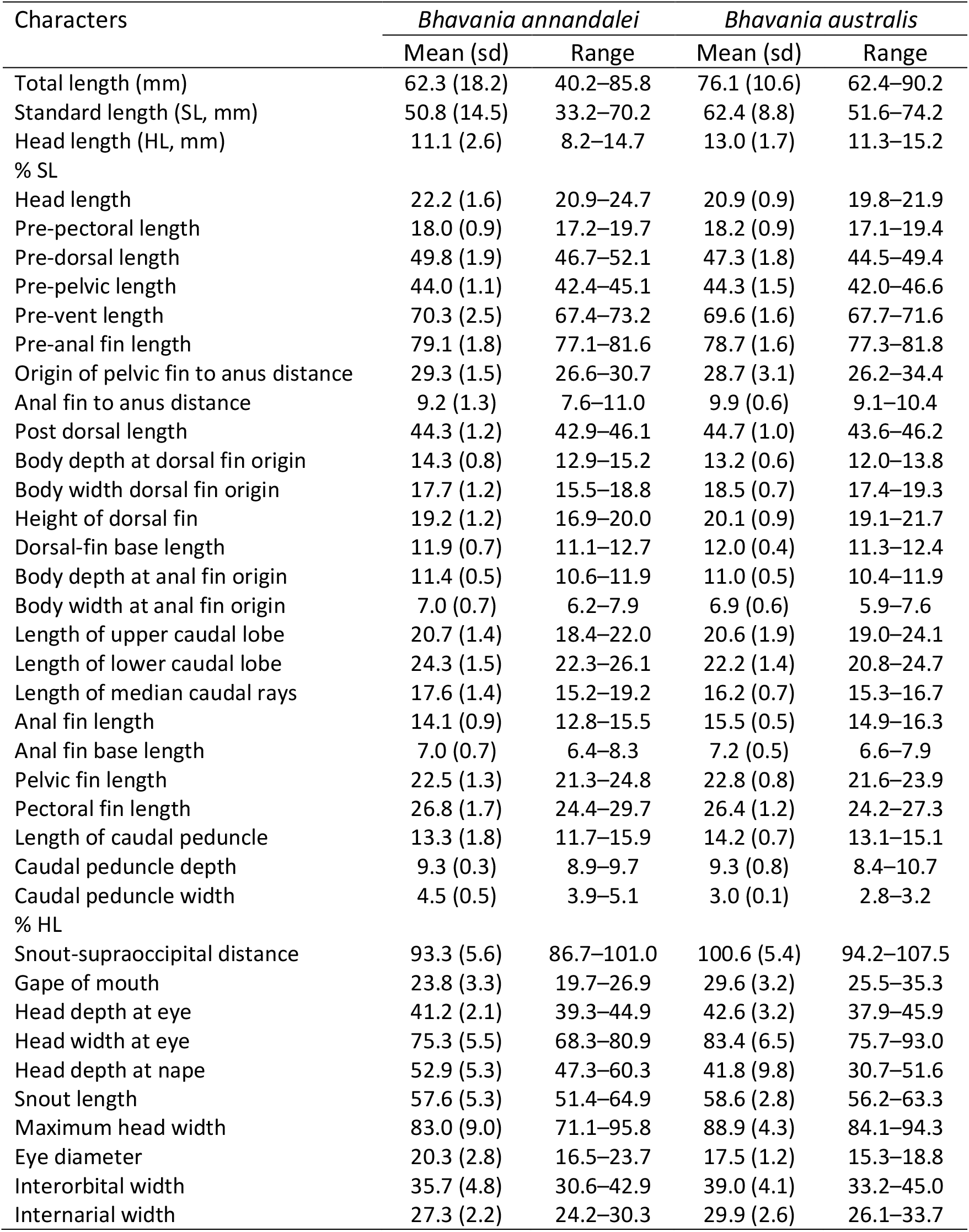
Morphometric data of *Bhavania annandalei* (KUFOS.19.AS.BH.02.1-6, n = 6) and *B. australis* (KUFOS.19.AS.BH.01.1-6, n = 6) putative topotypes.

**Table 2.** Meristic data of *Bhavania australis* (KUFOS.19.AS.BH.01.1-6, n=6), and *B. annandalei* (KUFOS.19.AS.BH.01.1-6, n=6) putative topotypes. Numbers in parenthesis indicate frequency of character state in the materials examined.

| Characters | *Bhavana annandalei* | *Bhavana australis* |
| --- | --- | --- |
| Pectoral-fin rays | 6+10+I (6) | 6+9+I (1), 6+10 (1), 6+10+I (4) |
| Dorsal-fin rays | ii+7+I (6) | ii+7 (3), ii+7+I (3) |
| Pelvic-fin rays | ii+7 (2), ii+8 (4) | ii+7 (4), ii+7+i (2) |
| Anal-fin rays | ii+5 (6) | ii+5 (4), ii+5+i (1); ii+6 (1) |
| Caudal-fin rays | 19 (6) | 19 (6) |
| Lateral line scales | 65+4 (2), 66+3 (1), 67+3 (2),<br>67+3 (1) | 65+3 (2), 65+4 (1), 66+3 (1), 68+3 (1),<br>69+3 (1) |
| Predorsal scales | 29 (1), 30 (2), 31 (3) | 28 (3), 29 (2), 30 (1) |
| Post dorsal scales | 34 (3), 35 (2), 36 (1) | 38 (1), 39 (2), 40 (2), 41 (1) |
| Scale above the lateral line | 11 (2), 12 (4) | 14 (4), 15 (2) |
| Scale below the lateral line | 9 (2), 10 (4) | 10 (3), 11 (3) |

## *Bhavania annandalei Hora* 1920

(Image 1, 2 and 3)

### Materials examined

KUFOS.19.AS.BH.02.1-6, 6 ex., 07.02.2019, 8.945 N & 77.158 E, 32.7–37.6 mm SL, Palaruvi falls, Tenmala, Kallada River, Kerala, India, coll. Arya Sidharthan, E.S. Abhijith, and George Joseph.

### Diagnosis

*Bhavania annandalei* is distinguished from its only known congener *B. australis* by a combination of characters including: low density and sparsely distributed tubercles on dorsal surface of head, especially on operculum, (vs. high density of tubercles on dorsal surface of head and operculum) (Image 3); eyes dorsally positioned (vs. dorso-laterally positioned) (Image 3); gape of mouth comparatively farther from snout tip, as a result the rostral barbels reaching anterior border of upper lip, (vs. gape of mouth closer to snout tip, and rostral barbels reaching posterior border of upper lip) (Image 3); rostral flaps between the rostral barbels fleshier (vs. less fleshier) (Image 3); fewer post dorsal scales (34–36 vs. 38–41); fewer scales above the lateral line (11–12 vs. 14–15); and caudal peduncle stout with its depth to width ratio 1.8–2.3 (vs. laterally compressed caudal peduncle with depth to width ratio 2.8–3.6).

### Description

Morphometric and meristic data are provided in Table 1 and Table 2, respectively. General body form as per Image 1a and Image 2a. Head details as in Image 3a, c.

Body elongate, depressed anteriorly, compressed laterally posteriorly; dorsal profile convex, deepest at dorsal-fin origin. Body wider than its depth at dorsal-fin origin, deeper than wide at anus. Head small, rounded, less than one-fourth of standard length; depressed, longer than broad, with minute sparsely distributed indistinct tubercles on dorsal surface of head. Eyes small, dorsally positioned, not visible from underside of head. Snout pointed in lateral view, round in dorsal view. Nostrils positioned dorso-laterally, closer to anterior border of eye than to snout tip, anterior nostril situated inside a skin flap covering the posterior nostrils. Mouth inferior. Lips fleshy. Gape of mouth less than three times maximum head width. Barbels three pairs, two rostral: outer rostral barbels shorter than inner ones; one pair of maxillary barbels, situated slightly anterior to the angle of mouth. Three fleshy rostral flaps interspaced between rostral barbels. Gill opening small, restricted above the base of the pectoral fin.

Body with scales except chest and belly. Lateral line complete, with 68–72 small scales. Caudal peduncle slender, its length almost three times its depth. Dorsal-fin originating slightly behind the pelvic-fin origin, closer to tip of snout than to caudal-fin base; with two unbranched followed by seven branched and a simple ray. Pectoral fin elongated, longer than head, with six unbranched, followed by ten branched and a simple ray. Pelvic-fin length almost equal to head length; fin origin closer to snout tip than to end of caudal peduncle, its posterior end not reaching anus, with two unbranched and eight branched rays. Anal fin with two unbranched and five branched rays. Caudal fin forked, with 19 principal rays.

### Colouration

In life (Image 1a), body is chestnut brown on dorsal and lateral sides, creamish-white on chest and belly; 3–4 prominent broad dark brown ventral bands; 2 broad ventral bands on the dorsal-fin base. There are 3 black-coloured bands on the dorsal fin, 6–7 bands on the pectoral, 3 bands on the pelvic, 1–2 bands on the anal, and 4 bands across the caudal fin.

### Morphometric analysis

In the morphometric analysis, using size-adjusted characters, the two species clustered separately on the first two PCA axes (Fig. 2a). The clusters were significantly different from each other (ANOSIM, 9999 permutations, R = 0.2315, P = 0.0271) indicating that the species formed distinct clusters in multivariate space. While length-length relationships for most characters showed similar trends for both the species, there were two relationships that showed marked differences. Length-length relationship between caudal peduncle depth and width (Fig. 2b) suggested that caudal peduncle width increased rapidly with increasing depth of the caudal peduncle in the case of *B. annandalei* as compared to *B. australis*. Similarly, length-length relationship between head length and head depth at nape (Fig. 2c) suggested that head depth increased rapidly with increasing head length in the case of *B. annandalei* as compared to *B. australis*.

**Figure 2.**
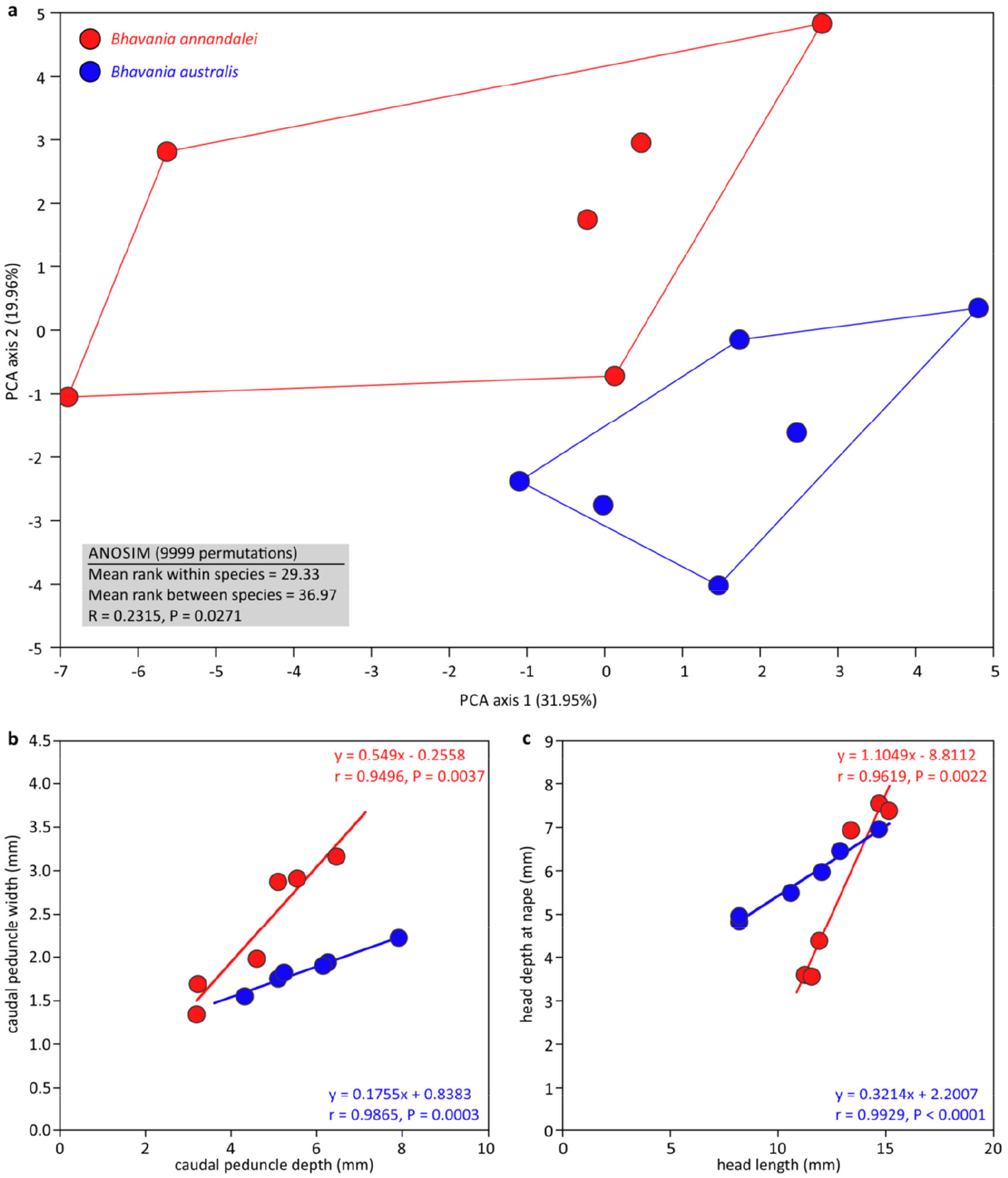
Morphometric analysis. (a) Principal Component Analysis scatter plot of factor scores and ANOSIM statistics. (b) Linear regression between caudal peduncle depth and width. (c) Linear regression between head length and head depth at nape.

### Genetic analysis

Topotypic *B. annandalei* (MT002520) differs from putative topotypic *B. australis* (MT002518) with a raw genetic distance of 6.4% in the *cox*1 gene.

### Distribution

*Bhavania annandalei* is known with certainty from the Kallada, Vamanapuram and Neyyar River systems in southern Kerala, India. These river systems drain the western slopes of the Agasthyamalai hill ranges, south of the Shencottah Gap. It is highly likely that the species also occurs on the eastern slopes of the Agasthyamalai hills particularly in the Tambaraparini River system in Tamil Nadu, but detailed surveys and voucher specimens are required to confirm this.

### Remarks

The density of chromatophores in *Bhavania* is likely to be dependent on the micro-habitat as well as the colour and type of substratum it inhabits. Other ecological factors that may influence body colour are forest/canopy cover, intensity of light, water flow and water temperature (V.K. Anoop Pers. Observ.). This is reflected in the different body colours shown by the two species in different habitats and locations (see Image 1), an observation which was also made by Hora (1941).

### Comparative material

*Bhavania australis*, KUFOS.19.AS.BH.01.1-6, 6ex., 13.04.2019, 10.8636 N & 76.6904 E, 46.4–58.8 mm SL, near Kavarakund falls, upstream of Malampuzha Reservoir, Kerala, India, coll. M.R. Ramprasanth.

## Acknowledgements

VKA and AS thanks the Kerala State Biodiversity Board (KSBB) for PhD fellowship, and RLS and RR thanks the Center for Aquatic Resource Management and Conservation (CARMAC), Kerala University of Fisheries and Ocean Studies (KUFOS) for funding. The authors are grateful to M.R. Ramprasanth, Josin Tharian, Vishnu Raj and Anvar Ali for useful discussions and help in the field. Permits for collection inside forest areas of Kerala were provided by the Kerala State Forest and Wildlife Department to VKA and AS.

---

**Image 1.**
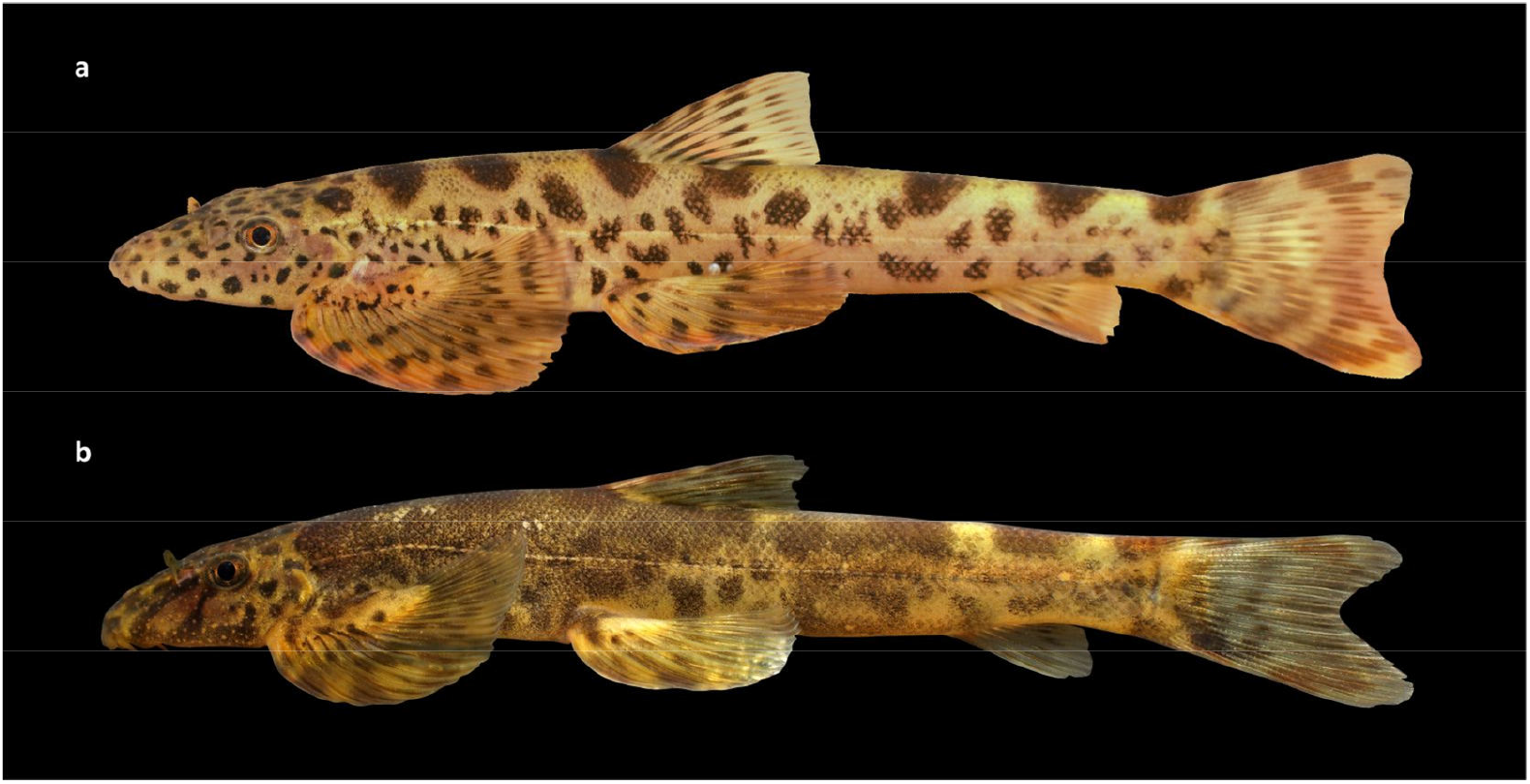
Putative topotypes of (a) *Bhavania annandalei*, and (b) *B. australis* in life (specimens not preserved).

**Image 2.**
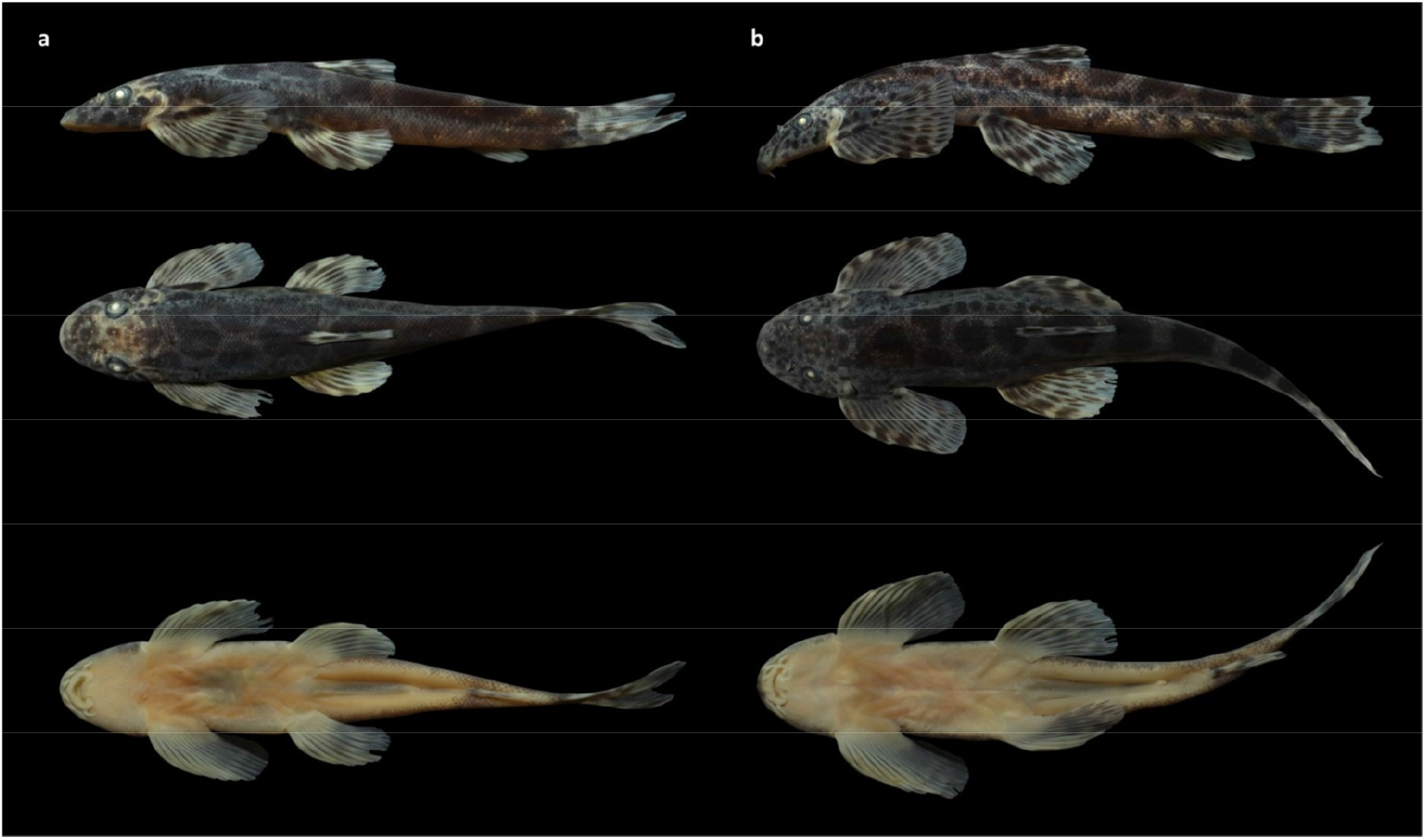
Dorsal, lateral and ventral images of putative topotypes of (a) *Bhavania annandalei* and (b) *B. australis*.

**Image 3.**
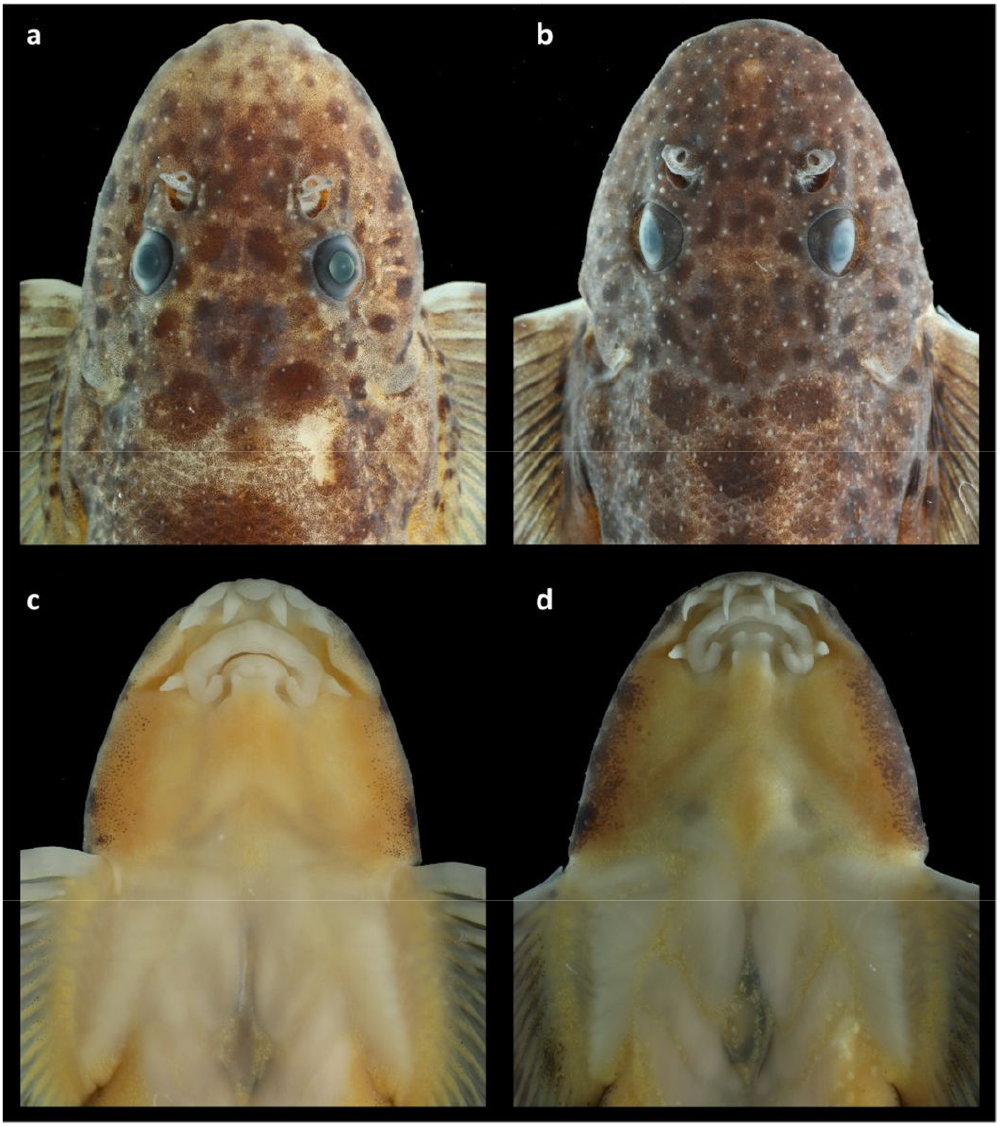
Dorsal and ventral view of head. (a, c) *Bhavania annandalei* and (b, d) *B. australis*.

